# Charged amino acid propensities and solubility: lysine is elevated at the termini of helices in *E. coli* proteins

**DOI:** 10.64898/2026.07.13.738247

**Authors:** Sifan Zhang, Jim Warwicker

## Abstract

An emerging result in the relationship between amino acid sequence and protein solubility is a preference, on average, for lysine over arginine in more soluble proteins. The termini of helices are known to be prone to partial unfolding, often employing N- and C-cap amino acids to maintain stability. Hypothesising that lysine/arginine differences in relation to solubility may be evident at helical termini, their propensities and predicted charge interactions in helices were examined. There is enrichment of lysine over arginine at helical termini in AlphaFold models of *Escherichia coli* proteins, more so (on average) for the most soluble proteins. Similar effects are seen for the sum of charged amino acids at helical termini. Regions other than helical termini also show correlation of lysine composition, and overall charged amino acid composition, with solubility. These results suggest that protein design protocols could improve solubility through targeting lysine enrichment in regions such as helical termini, in addition to the more conventional consideration of helix capping interactions.

## 1. Introduction

Protein solubility, the measure of how much protein can dissolve in solution, is crucial for biological function in aqueous systems. This importance relates to physiological and biotechnological scenarios. Indeed, expression of protein at high yields has been a major driving force for increasing understanding of protein solubility, and deriving predictive models based on amino acid sequence and/or structure. The endogenous cell cytoplasm appears to be constituted such that solubility of a protein is tuned to expression level, from observation of an inverse relationship between expression (mRNA) level and aggregation rate for *E. coli* proteins [1]. It is therefore unsurprising that engineering an organism for increased production of an exogenous or endogenous protein can lead to issues with solubility and aggregation. Such problems are common in recombinant protein production and are increasingly becoming evident as a limiting factor in engineered synthetic biology pathways [2].

Stability and solubility are closely linked properties for proteins. The prominent Lumry– Eyring model for irreversible protein aggregation invokes a non-native aggregation-prone, intermediate [3]. Such intermediates may include solvent exposed non-polar surfaces. It is well-known that pH modulates protein solubility via net charge. Thus, charge and hydrophobicity each play a major role in modulating protein solubility, as demonstrated in sequence-based machine-learning models for biophysical properties of clinical-stage antibodies [4].

Alongside the design of protein stability, itself undergoing a renewal with machine-learning models for sequence relationships to structure [5], protein solubility prediction has become a fertile area. However, data availability is lower for protein solubility as compared to protein structure. The latter case includes sequence to structure relationships encoded in multiple sequence alignments, that (generally) imply amino acids proximity in 3-dimensions. The largest datasets for protein solubility have arisen from various programmes for high-throughput protein production, typically structural genomics in the early 2000s, and imposition of passage through the purification pathway at some stage as indicating more soluble protein, with a similar rationale applied for selecting proteins from the protein structural database (RCSB [6]) [7]. In contrast to these very coarse (binary) assessments of protein solubility, high-throughput quantitative experimental data are very limited. The most widely used dataset is that derived from cell-free expression of *E. coli* proteins [8]. One of the earlier sequence-based models to be published, recapitulated the importance of charge and hydrophobicity in fitting to the experimental data, but also discovered a relationship between lysine and arginine composition and solubility [9]. The model was subsequently delivered as a web tool [10]. More soluble proteins, on average, prefer lysine over arginine in the *E. coli* proteins, leading to the suggestion that the difference between lysine and arginine sequence compositions (termed [K – R]) could be a design tool for improving protein solubility. This was borne out experimentally in swaps of arginine and lysine in a single chain Fv fragment [11].

Constructing predictive models for protein solubility is a rich area of research, although with limitations imposed by the lack of high-quality datasets [12]. Models that incorporate protein structure have been reported [13–15], along with those directed at predicting the solubility of protein mutants [16, 17]. Prominent in recent years have been models based on machine learning [18]. Alongside developments in predictive models and the requirement for increased collection of high-quality datasets, there remain questions of biophysical understanding i.e. how sequence and structure couple to mediate protein insolubility. A prime example of this is the observation that lysine is preferred to arginine, on average, in the more soluble proteins of *E. coli*.

The lysine/arginine effect couples with the relative importance of protein abundance. Less abundant proteins, that do not contribute as much to the cellular protein content, do not exhibit the preference for lysine. Indeed, protein solubility is correlated with protein abundance for *E. coli* [10]. Current understanding for lysine to arginine difference with regard to solubility is that arginine is more strongly interacting in different structural environments, from salt-bridge interactions [19] to cation-pi interactions [20]. There follows the possibility that arginine could mediate interactions with aromatic-containing regions from partially unfoldeded segments, similar to the role of cation-pi interactions in FUS phase separation [21], and thus that arginine could be more prone (than lysine) to interactions that promote formation of aggregate nuclei.

A prime candidate for regions of increased flexibility and partial unfolding propensity in structured regions of proteins are the ends of alpha helices, evidenced by NMR-based study of flexibility [22], and stabilising motifs at helical termini [23], including N-cap and C-cap amino acids [24]. Given the potential for transient unfolding and a concomitant increased exposure of non-polar surface at helical termini, it is hypothesised that there could be a compensating elevation of lysine relative to arginine composition, particularly in more soluble (and abundant) proteins. The current work studies the distributions of amino acids along α-helices, concluding that there is indeed variation in more soluble proteins, suggesting a potential design feature for solubility to mimic occurrence in naturally occurring proteins.

## 2. Materials and Methods

### 2.1 Proteome sequence and structure modelling

AlphaFold version 2 protomer models were obtained for 4463 *E. coli* proteins from the AlphaFold Protein Structure Database hosted by the European Bioinformatics Institute (EBI, alphafold.ebi.ac.uk, [25]), matching the proteome sequences file UP000000625_83333.fasta from UniProt [26]. Proteins with predicted transmembrane segments were excluded from analyses, using Kyte-Doolittle predictions of hydropathy [27], and a threshold of 1.6 (over 21 amino acid windows), derived from protein-sol calculations [10]. The pLDDT (predicted local distance difference test) measure for AlphaFold models was used to assess reliability of predicted structure. A pLDDT threshold of 90 was used (from the 0 to 100 scale) to ensure that secondary structure analysis was restricted to high confidence predictions of secondary structural elements.

### 2.2 Secondary structure assignment, helical termini distances, and amino acid distributions in helices

Secondary structure was assigned with DSSP (Define Secondary Structure of Proteins) software [28], [29], concentrating on assignment of α-helices. For the purposes of calculations as a function of distance from helical termini, distances (within a helix) from helical N-terminus (NT) and C-terminus (CT) were calculated, together with the shorter of these (SH). As amino compositions are sampled for increasing distances from helical data, the number of data points decreases. An estimate of the standard deviation for each location-amino acid type pair was taken as the square root of the instances of that pair, and a threshold set for NT, CT, SH, such that the standard deviation remains below a certain value for each analysis, as given in the Figure Legends.

### 2.3 Calculation of charge interactions

Continuum electrostatics methods were used to predict ΔpKa values for protomer AlphaFold proteins in the *E. coli* proteome. Structure-based calculations were made with locally installed code for solvent accessibility surface area (SASA, sacalc), and for pKa (pkcalc, PROPKA3) [30, 31]. A threshold magnitude ΔpKa of 2 was applied for these calculations and values above this were omitted. For some data analysis ΔpKas were transformed for more convenient visualisation, such that positive indicates stabilisation of the ionised state relative to the unionised state. A set of calculations without helix dipole (peptide dipole) charges was made with pkcalc by removing the amino acid main-chain charges from the dictionary. Additional calculations used a simple Debye-Hückel prediction of ΔpKas, with a single continuum relative dielectric of 80, and a continuum monovalent salt concentration of 0.15 M. PROPKA3 and pkcalc also gave pH-dependent folding stability, for which the value at pH 7 was combined with protein length to give an estimate of the ionisable charge energy per amino acid. A distribution of experimental pKas, across organisms, was obtained from the PKAD-R database (https://sites.krieger.jhu.edu/damjanovic-lab/pkad-r/ [32]). Calculation of protrusion at a protein surface was made by implementing a protocol for the Cx method [33].

### 2.4 Protein solubility data

Experimental solubility data for 3049 *E. coli* proteins [8] were filtered to remove those containing a predicted transmembrane TM region with Kyte-Doolittle hydropathy > 1.6 [27], leaving 2385 data points.

## 3. Results and Discussion

Alpha-helices in AlphaFold protomer models for the *E. coli* proteome [26, 34] were annotated with secondary structure (DSSP [29]). Helices with predicted transmembrane (TM) segments from Kyte-Doolittle [27] sequence analysis [10] were excluded, to focus on water soluble proteins. To ensure confidently predicted helices in the native state, only regions with pLDDT ≥ 90 were included.

### 3.1 Amino acid distributions along helices of the *E. coli* proteome

The distributions of amino acids (AAs) throughout helices, as percentage deviations from the average for each amino acid (average over locations from 1 to 15 from helical termini) were calculated (Fig 1). Clear signals are seen for Asp and Glu at helical termini in the heat map (denoted SH) that combines distances to N- and C-termini (NT and CT, Fig 1A). This resolves into the expected N-terminal and C-terminal interaction preferences when viewed in the equivalent heat maps for separated NT and CT distances (Fig 1B/C). There is variation, dependent on location, with the precise N-cap position excluded since it lies before N1 (the first AA with mainchain angles with alpha helix conformation), but results are consistent with measured AA preferences at N1 and N2 positions [24, 35, 36]. There are also the anticipated signals (enrichment in the SH plot) for Lys and Arg that resolve to the CT (Fig 1B/C), with a small signal for His. Some polar and uncharged AAs (Q more so, N and S somewhat) are also enriched at termini (with variation between NT and CT). Proline is enriched at the NT, but not at CT helical termini, and is the most highly depleted AA within helices, consistent with its role as a helix breaking amino acid [37]. Glycine is depleted towards the CT. Perhaps reduced Gly lowers flexibility in regions that could destabilise a protein with partial unfolding. AAs with hydrophobic sidechains (A, V, I, L, F, Y) are also depleted near helical termini, to varying degrees, presumably due to increased solvent exposure in these regions. There is also an overall periodic distribution for most AAs, opposite for charged/polar and hydrophobic amino acids, matching the solvent accessible surface area (SASA, Fig 1D), consistent with the differences in AA solubilities.

**Fig 1.**
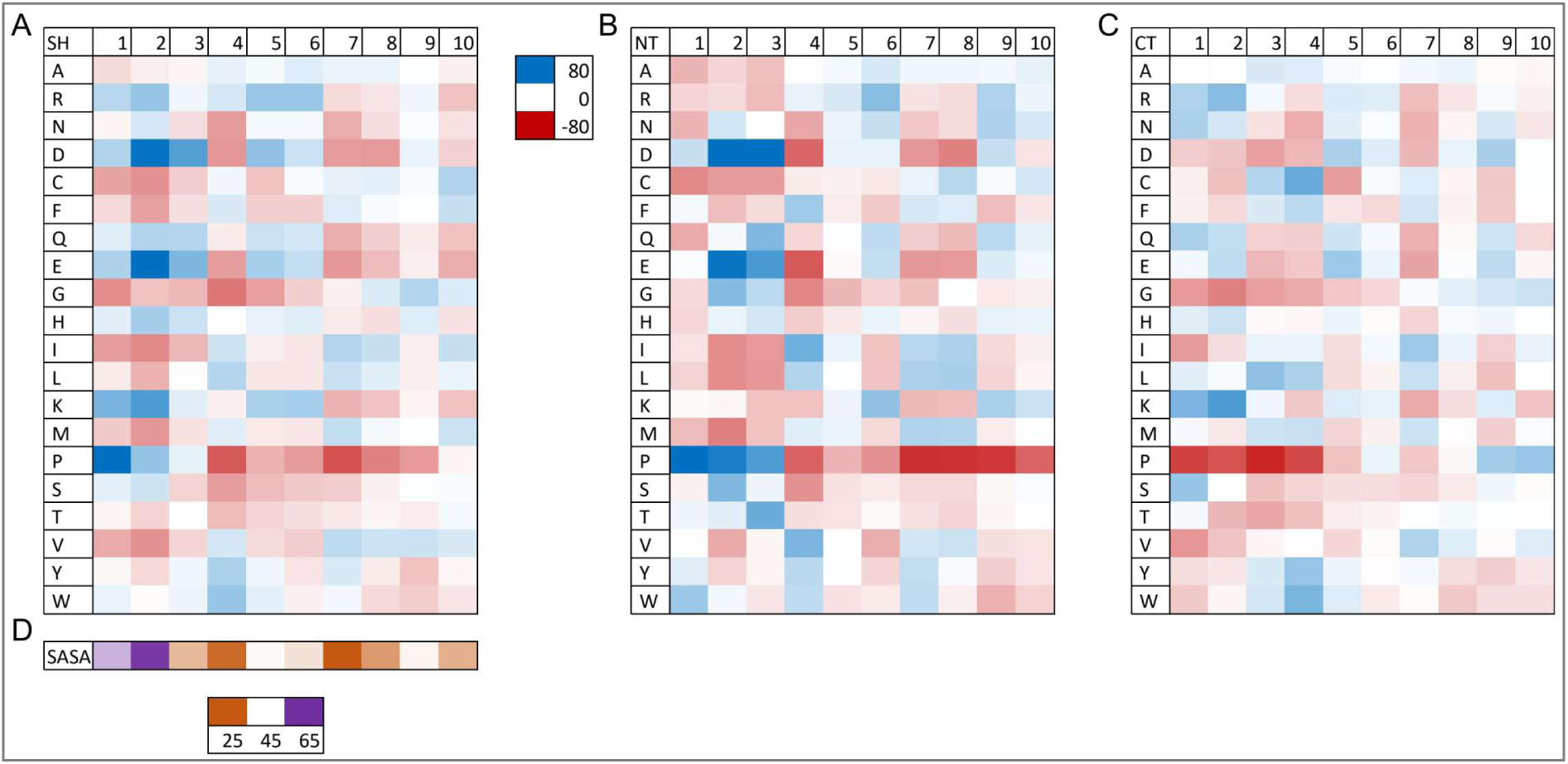
Amino acid propensities in well-defined helices of *E. coli* proteins. In panels A, B, C, for locations from 1 to 10 relative to helical termini, amino acid percentage deviations from the average for that amino acid, calculated over locations from 1 to 15 relative to helical termini, are displayed in heat maps (from red -80% to blue +80%). Single letter amino acid codes are shown in the left-most column. (A) Shortest distance to a helix terminus, (B) distance to N-terminus, (C) distance to C-terminus. (D) Average SASA (Å^2^) over AAs for shortest distance to a helical terminus, matched with the locations in panel (A), facilitating comparison of site burial/exposure and AA propensities.

It is apparent that variation of charged and polar AA enrichment, and hydrophobic AA depletion, near helical termini, is extensive. Beyond the clear coupling to SASA (e.g. avoiding the burial of charged AAs), it is of interest to investigate charged AA compositions and predicted charge interaction energetics at helical termini, beyond the helix cap positions.

### 3.2 Helices in the most soluble *E. coli* proteins have increased charge composition towards their termini, with lysine favoured over arginine

AAs that normally carry a net charge at neutral (close to cytosolic) pH are Lys, Arg, Asp, and Glu. Summed percentage compositions for these AAs at each helical location (measured as SH, NT and CT distances) show some enrichment towards termini, position 2 in particular (Fig 2A). When viewed for only the 500 most soluble proteins (Fig 2A), this enrichment is greatly enhanced, and the expected periodicity (correlating with ASA, Fig 1) is evident. The overall increased prevalence of charged amino acids at helical termini will in part result from increased SASA due to the absence of peptide-peptide hydrogen-bonding. A large increase in the prevalence of summed K, R, D, E for the most soluble *E. coli* proteins is consistent with evolution towards an increase in stability at helical termini for these proteins. Since this enrichment extends beyond the first helical turn (Fig 1A), it does not arise solely from increased SASA at the first turn of each helical terminus. A parallel analysis was made for the net charge from the difference of (Arg + Lys) and (Asp + Glu) compositions (Fig 2B). The most dominant feature is an enrichment of negative charge close to the NT, as expected, with a smaller enrichment of positive charge near to the CT. There is some increase in the enrichment of negative charge for the set of 500 most soluble proteins, compared with the whole protein set.

**Fig 2.**
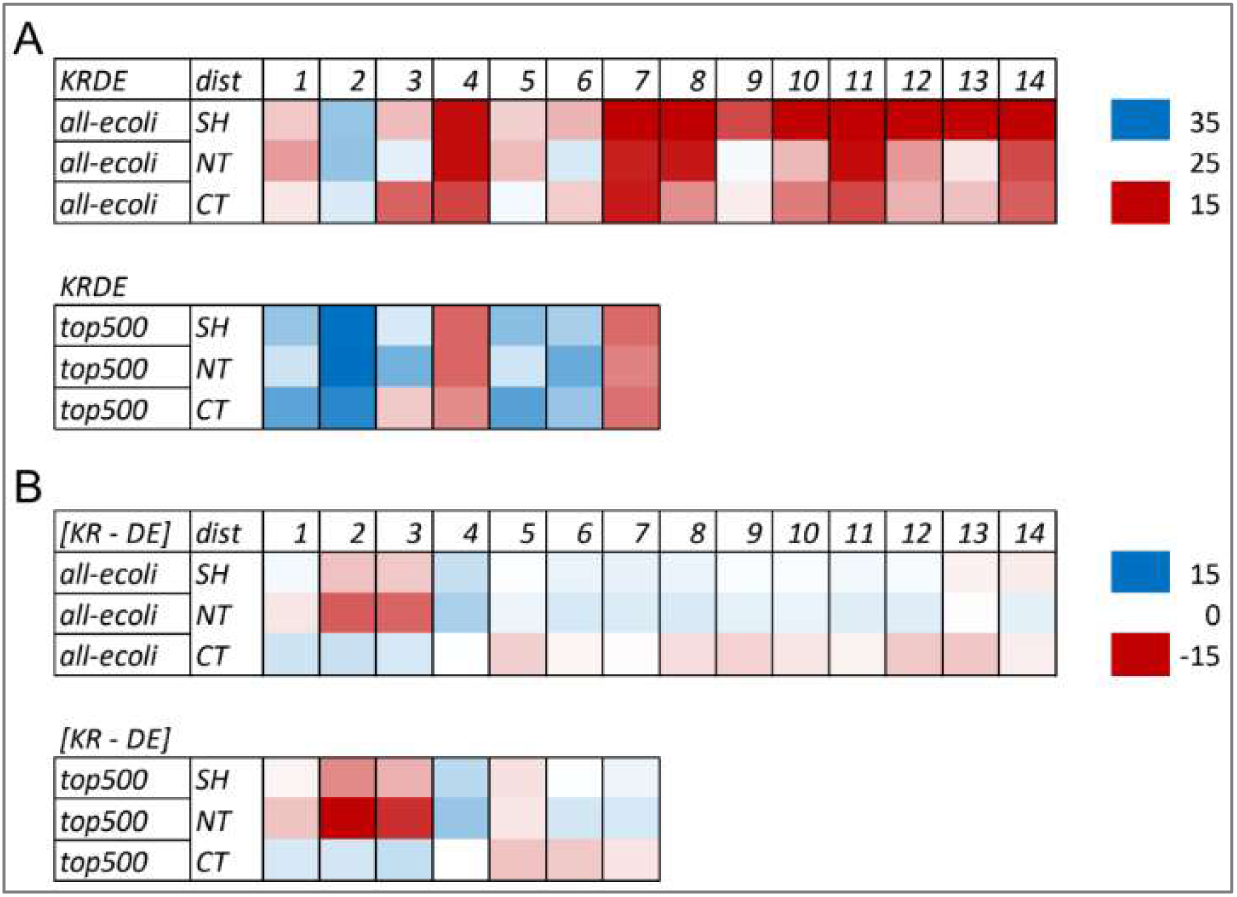
Charged amino acids are enriched at helical termini. (A) The summed percentages for K, R, D, E (out of all AAs at that location) are displayed as a heat map (from red, 15% through white, 25% to blue, 35%). Position within a helix is numbered from N-terminus (NT), C-terminus (CT), and the termini combined (shortest distance to terminus, SH). The upper panel is for all helices that meet the 90% pLDDT threshold and lack a predicted TM segment, with the locations 1 to 14 shown, for which the estimated standard deviation < 1% (Materials and Methods). Using the same criteria for helices from only the 500 proteins with the highest measured solubility [8] (lower panel), give reliable data from positions 1 to 7. (B) Heat maps for the difference between summed K, R and summed D, E (the net charge, assuming minimal contribution from other AAs). Here the colours are such that red is negative and blue positive. Upper panel for all proteins, lower panel for the 500 most soluble proteins.

The difference in Lys and Arg composition [K – R] of a protein sequence is one of the most significantly correlating properties with *E. coli* protein solubility [10]. Since charged AAs are enriched at helical termini, and particularly in the most soluble proteins, [K – R] was also investigated in these regions. Arg is generally at higher percentage than Lys in helices (Fig 3A, upper panel), but this difference falls towards the termini. When only the 500 most soluble proteins are included, then Lys composition increases relative to Arg. As an example of the numbers, for the shortest distance (SH) to a helix terminus of 2, Lys and Arg percentages are 5.78 and 7.02 for the larger set, with 7.39 and 6.62 for the most soluble protein set. In the equivalent analysis, with [K – R] replaced by [D – E] (Fig 3B), an overall preference for E over D in helices is apparent, and somewhat greater in the most soluble protein set. Despite this substantial preference for E over D at helical termini, the correlation of [D – E] with solubility is much lower than that of [K – R] (Table 1).

**Fig 3.**
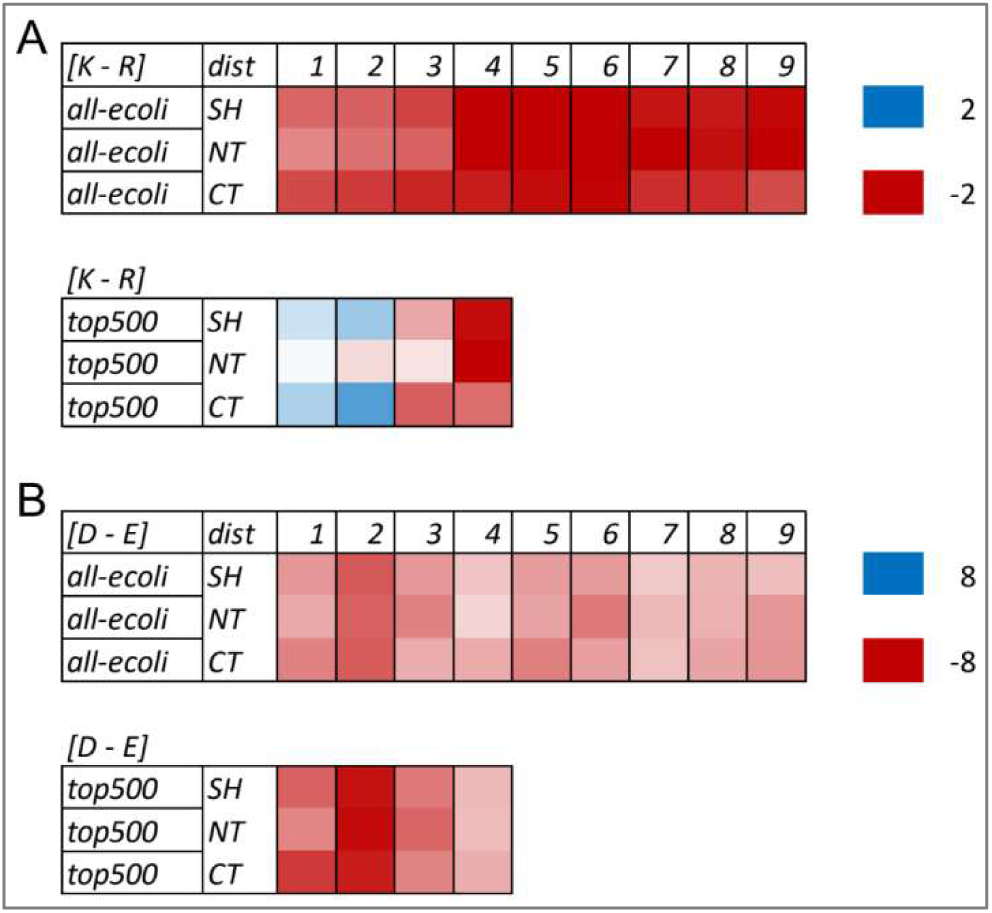
Lysine is enriched relative to arginine at the termini of helices in the most soluble *E. coli* proteins. (A) The difference in percentage amino acid compositions of Lys and Arg [K – R] is shown versus location in all *E. coli* protein helices that meet the 90% pLDDT threshold and lack a predicted TM segment (upper panel). The heat map runs from red, -2% through white, 0% to blue 2%, and displayed for locations 1 to 9 for which the estimated standard deviation < 0.3%. Equivalent data are shown for helices in the 500 most soluble proteins, here with estimated standard deviation < 0.44%, giving locations 1 to 4. (B) Equivalent data are shown for the difference in percentage compositions of Asp and Glu [D – E], all proteins (upper panel) and the 500 most soluble proteins (lower panel). For [D – E] data, the heat map runs from -8% (red) through 0% (white) to 8% (blue).

**Table 1.** Percentage compositions and correlations with solubility, for combinations of charged AAs in *E. coli* AlphaFold protomer models.

footnotes: Percent composition for the listed amino acid combinations, together with correlation *versus* experimental solubility for those combinations, for *E.
|  | percent | percent | percent | percent | corr-solub | corr-solub | corr-solub | corr-solub |
| --- | --- | --- | --- | --- | --- | --- | --- | --- |
|  | KRDE | [KR – DE] | [K – R] | [D – E] | KRDE | [KR – DE] | [K – R] | [D – E] |
| all AAs | 23.11 | -1.48 | -0.92 | -0.98 | 0.292 | -0.113 | 0.289 | -0.108 |
| all-ge90 | 22.69 | -1.56 | -1.13 | -1.05 | 0.235 | -0.079 | 0.237 | -0.069 |
| hel-ge90 | 25.35 | -0.77 | -1.56 | -3.86 | 0.125 | -0.08 | 0.159 | -0.078 |
| all-lt90 | 24.24 | -1.24 | 0.29 | -0.62 | 0.071 | -0.031 | 0.124 | -0.026 |

### 3.3 Lysine to arginine ratio varies throughout *E. coli* proteins

The property [K – R] features in the protein-sol model for protein solubility due to a correlation of 0.289 with measured solubility [10], comparable with other highly correlating features (KRDE / 0.292, FWY / -0.263, protein length / -0.367). Table 1 gives percentage compositions and correlations with solubility for combinations of charged AAs. The combinations KRDE and [K – R] have substantially higher correlation with solubility than [KR – DE] and [D – E]. Correlation values of KRDE and [K – R] with solubility are not higher for any of the AA categories in Table 1 than that with all AAs. This is consistent with a systematic difference of [K – R] between the categories, as well as within them. For example, [K – R] percentage composition, excluding proteins with predicted TM segments, is -1.56 for helical AAs with pLDDT ≥ 0.90 and 0.29 for all AAs with pLDDT < 90. Correlations with solubility for those two subsets of AAs are 0.159 and 0.124, respectively, indicating that it is not just the most well-defined structural regions for which Lys relative to Arg composition is relevant to protein solubility. A possible molecular mechanism for reducing arginine-mediated cation-pi interactions between partially unfolded proteins, suggested for helical termini, may extend to other environments [38]. It is interesting that while the correlation of FWY (summed aromatic AA compositions) with solubility is -0.263 for all AAs in proteins lacking a predicted TM segment, that for calculated Kyte-Doolittle values (representing overall hydrophobicity) and solubility is of much small magnitude at -0.103, consistent with the hypothesis of Arg and aromatic residues coupled in contributing to protein insolubility.

### 3.4 Distributions of predicted charge interactions show a small but systematic difference between calculation methods

To predict charge-charge interactions in *E. coli* protomer AlphaFold models, two methods were compared, the commonly used PROPKA3 [31] and the Finite Difference Poisson-Boltzmann (FDPB) and Debye-Hückel (DH) combined method pkcalc [30]. These two packages were also used to analyse charge interactions in human AlphaFold protomer models [39]. While PROPKA3 and pkcalc both give peaks in the distributions for stabilising ΔpKas between 0 and 1, the range expected for favourably interacting, solvent exposed groups [40], the PROPKA3 distribution also has a large contribution in the weakly destabilising region (Fig 4A). These predictions for *E. coli* proteins were compared with a set of experimental pKas for proteins across species from the PKAD-R database [32] (Fig 4B). The experimental data were adjusted to ΔpKas using either PROPKA3 or pkcalc intrinsic pKa values for D, E, K, R. There is not a large difference in those adjusteddistributions (Fig 4B). The mildly destabilising region is not as populated in the experimentally derived distribution as in the PROPKA3 predictions. This is not to say that PROPKA3 predictions are necessarily worse overall than pkcalc. In a previous study PROPKA3 was superior in ΔpKa-based prediction of groups that mediate functional pH-dependence, where pkcalc can over-predict ΔpKas in less solvent accessible regions [39]. The combination of FDPB and DH protocols in pkcalc ameliorate this issue to some extent, without more computationally expensive conformational sampling techniques. There may also be some bias in the experimental set, in that 6% of the ΔpKas are < -2 (and excluded), possibly due to a degree of focus on measurements for destabilised, typically enzyme active site, groups. With regard to the peak of ΔpKa values at a mildly destabilising level, it is possible that the heuristics-based desolvation energy calculation in PROPKA3 slightly overestimates the burial penalty for some mostly solvent-exposed ionisable groups.

**Fig 4.**
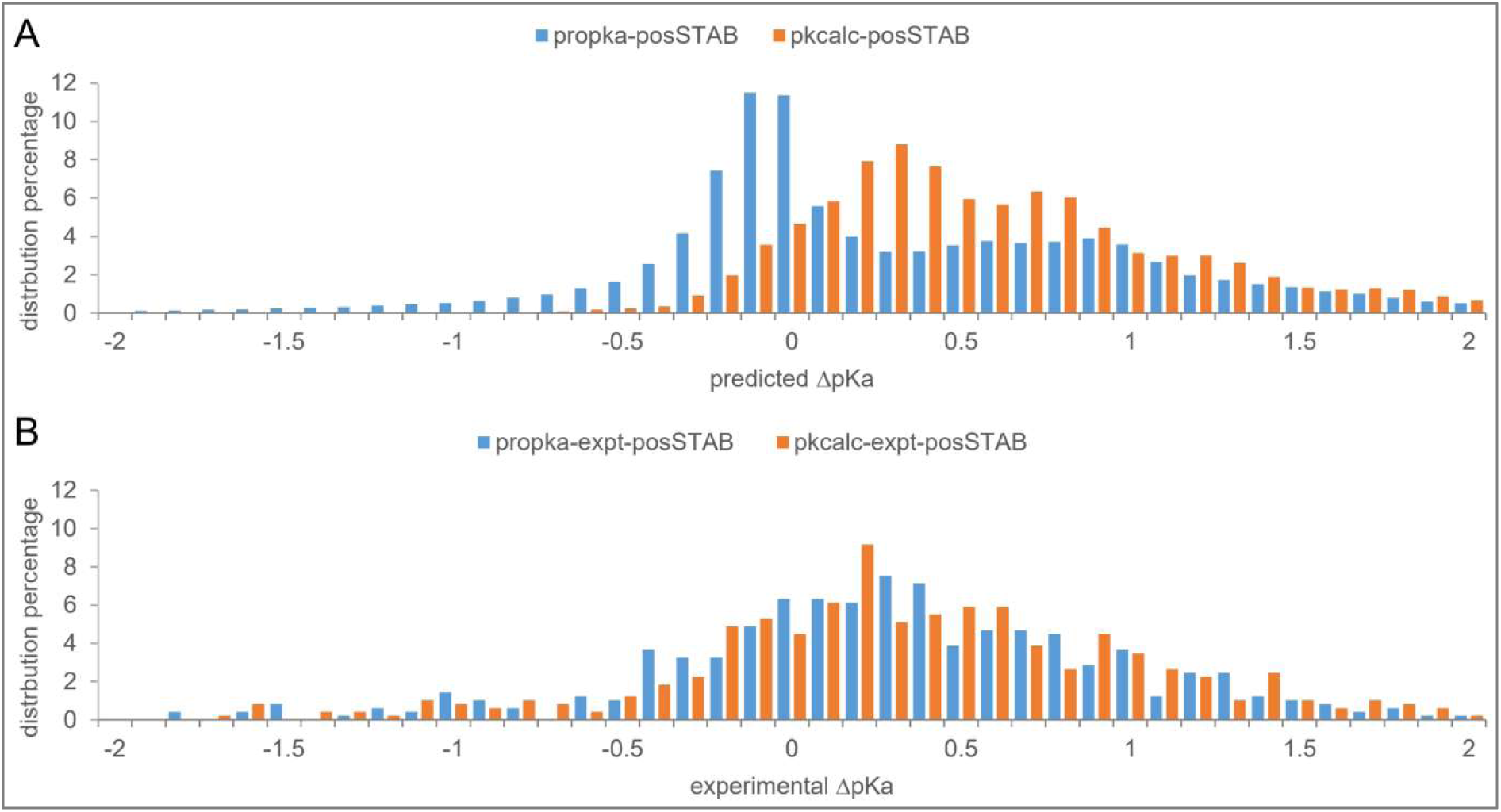
Comparison of ΔpKa calculations for *E. coli* protomer models. Distributions of calculated ΔpKas are shown for PROPKA3 and pkcalc methods applied to AlphaFold protomer models, with all results presented as positive ΔpKa relating to stabilisation of the ionised form (A). An experimental database of pKas from proteins across species are adjusted to ΔpKa according to either the intrinsic (model compound) pKa values used by propka3 or pkcalc, and displayed such that the range can be compared with the predicted ranges for *E. coli* protein ΔpKas (B). Measured and predicted ΔpKas with magnitude > 0.2 are excluded from the analysis.

Both prediction methods show distributions for ionisable group types that largely follow their combined plots, with D (for both methods) having the highest percentage contributions of groups with higher stabilising ΔpKas (Fig 5). The discrepancy at mildly destabilising ΔpKa is present across all 4 AA classes.

### 3.5 There is no substantial increase of charge interactions towards helical termini

Charge interactions along helices were studied with ΔpKas averaged over groups (D, E, K, R separately) at each shortest distance to helical terminus location, for PROPKA3 and pkcalc predictions (Fig 6). Other than larger interactions at a distance to terminus of 3 AAs, there is no gradient of predicted interaction scale along helices. Thus helical termini mostly do not appear to be stabilised in terms of charge interactions at each group, although the composition of DEKR does increase towards the termini (Fig 2A). As expected from the overall distributions (Figs 4, 5), predicted PROPKA3 stabilisations (Fig 6A) are lower than those from pkcalc (Fig 6B). Predictions with pkcalc were repeated for AlphaFold protomer models with peptide dipoles carrying no charge, thus removing helix dipole interactions with amino acid sidechains (Fig 6C). There is some decrease of interaction over the first 3 locations from the closest helical terminus, the largest reduction at position 3. Interestingly, charged amino acids at the N3 position have been reported to impact helicity [41]. Overall, in terms of charge properties studied in the current work, that may be considered in addition to N-cap and C-cap stabilising interactions, there is some increased composition of KRDE towards termini, and a small additional stabilisation around position 3, that may be associated with mainchain charge interactions.

### 3.6 Protein length correlation with insolubility is a key and potentially multifaceted feature

Both PROPKA3 and pkcalc give predicted pH-dependent stability data, from which the predicted stability of folded relative to unfolded states was extracted at pH 7, for *E. coli* AlphaFold protomers. It was hypothesised that values of predicted ionisable group mediated stability (scaled for protein length), may correlate with solubility, via increased conformational stability. There is, however, only small a correlation for pkcalc and a strong anti-correlation for PROPKA3 (Fig 7A). Hypothesising that the latter result is associated with the peak of destabilising predicted ΔpKas for PROPKA3, a third calculation was made. The dhcalc method uses just Debye-Hückel calculations of charge interactions, with no contribution from desolvation, which is generally the biggest contribution to destabilising ΔpKa. Indeed, dhcalc gives a positive (although weak) correlation with solubility (Fig 7A). This result is expected from comparison with a database of experimental ΔpKa values (Fig 5), and a general understanding since early work [42] that surface charges are arranged to, on average, stabilise folded proteins. It seems unlikely that a large proportion of proteins would not be stabilised by ionisable group interactions at pH 7 (Fig 7B), despite the intriguing correlation of PROPKA3 predicted stability with solubility (Fig 7C).

**Fig 5.**
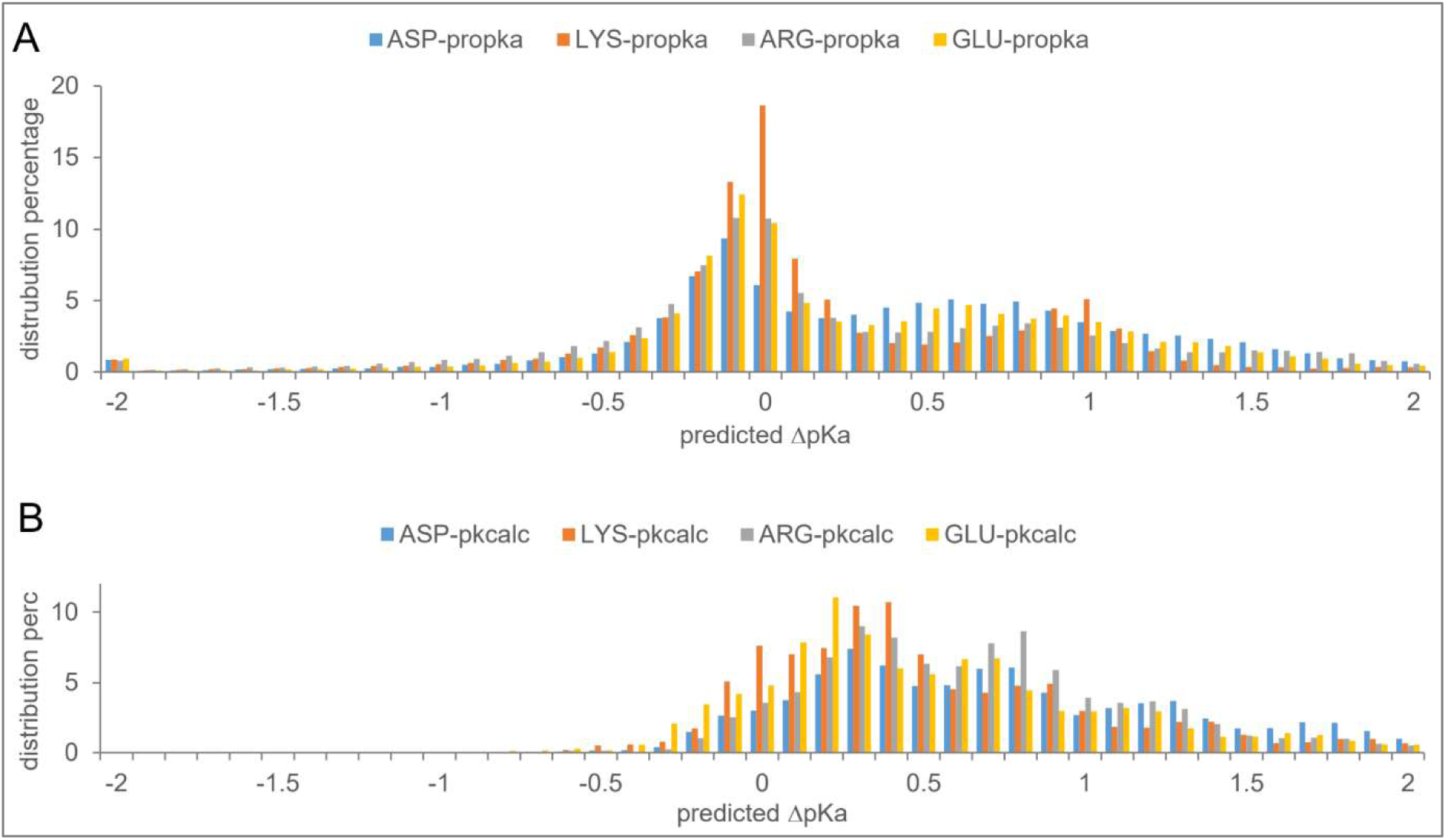
Amino acids follow similar ΔpKa distributions. Distributions of calculated ΔpKas are shown for each for the D (ASP), E (GLU), K (LYS), R (ARG) AA sidechains ionisable groups, PROPKA3 (A) and pkcalc (B). ΔpKa values have been adjusted such that positive relates to stabilisation of the ionised form.

**Fig 6.**
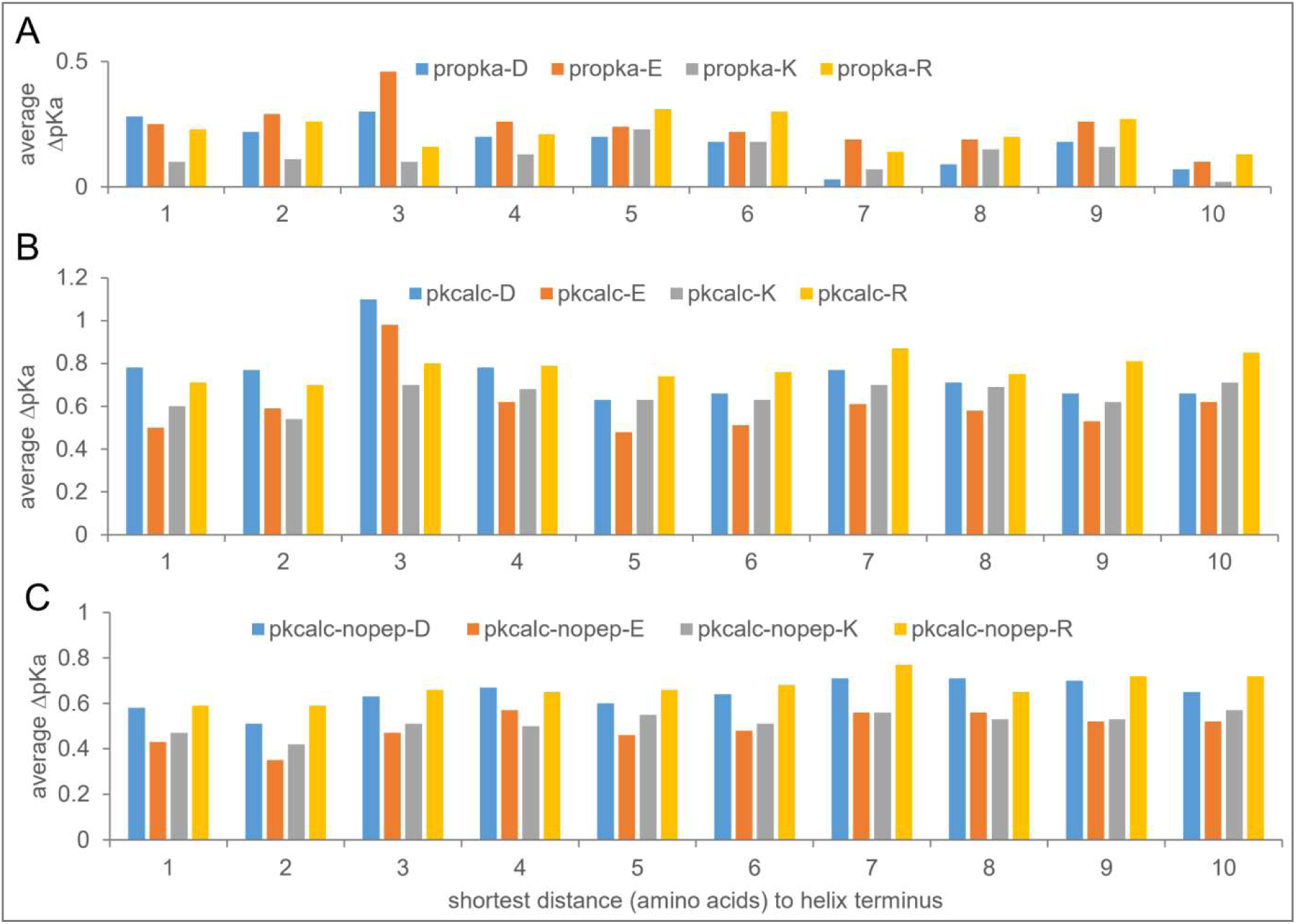
Predicted charge interactions are relatively uniform along a helix. ΔpKas for helical AAs, calculated with PROPKA3 (A) or pkcalc (B), are shown for the first 10 AAs to the nearest helix terminus. Results are separated by ionisable sidechain type, all values are averaged over AAs at that position, with positive ΔpKa reflecting stabilising interactions. (C) Calculations with pkcalc are repeated for systems where the peptide dipole charges of the protein mainchain are zero.

**Fig 7.**
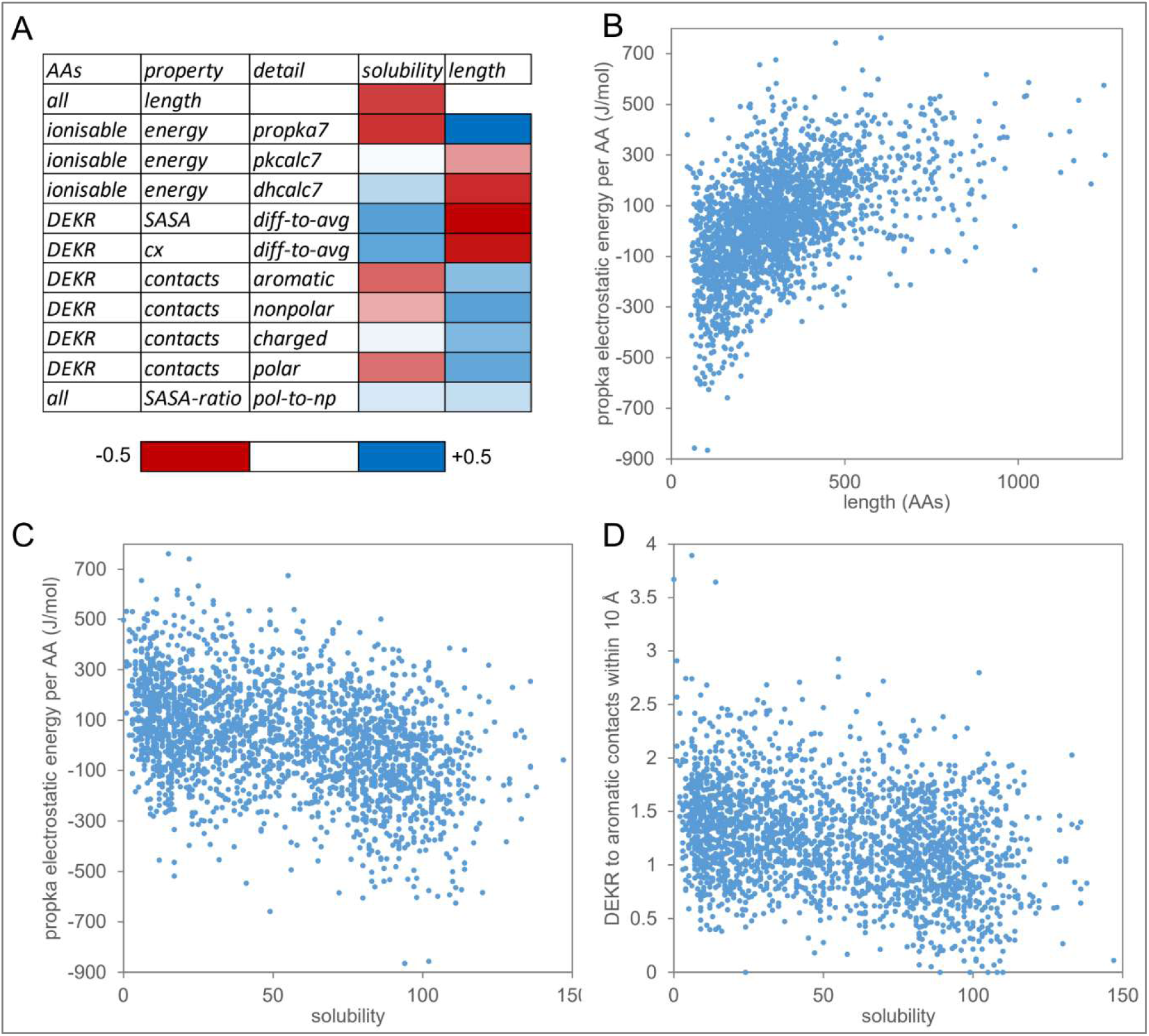
Negative correlation of solubility with protein length associates with geometry at the protein surface. (A) Correlations between either experimental solubility or length of *E. coli* proteins and various calculated properties are shown in a heat map, from negative (-0.5) to positive (+0.5). Properties include calculated sum of ionisable group stabilities at pH 7 from PROPKA3 and pkcalc (normalised for protein length); SASA and Cx (protrusion) values for D, E, K, R relative to averages over the *E. coli* proteome; contacts to D, E, K, R groups from different AA classes; ratio of polar to non-polar SASA for all AAs. (B) Scatter plot of PROPKA3 summed ionisable group energy (normalised for protein length, per AA, J/mol), against protein length. (C) Scatter plot of PROPKA3 summed ionisable group energy (per AA, J/mol) against experimental solubility. (D) Scatter plot of average contacts to aromatic AAs to D, E, K, R AAs (within 10 Å), against experimental solubility.

It is interesting that predicted PROPKA3 stability correlates with solubility, with correlation coefficient of similar size to the largest contributing features to the protein-sol model. A likely resolution, in part, is the anti-correlation of PROPKA3 stability with protein length, which itself anti-correlates with solubility and is in the protein-sol model. The question remains of why PROPKA3 energy, scaled to a per AA value, is related to length. Considering that geometrical input to PROPKA3 could be leading to desolvation terms responsible for the peak in destabilising ΔpKas and the correlations with length and solubility, various features were calculated for D, E, K, R charged groups (Fig 7A). Solvent accessibility and a protrusion measure (Cx, [33]), also strongly anti-correlate with length, and correlate with solubility (Fig 7A). It is possible that the PROPKA3 desolvation term relates to the SASA and Cx changes with protein length, where desolvation penalty increases with protein length. The differences in predicted ΔpKa distributions for PROPKA3 and pkcalc indicate that such effects are much smaller for the FDPB-DH method of pkcalc. This geometrical reasoning coupled to an over-estimate of desolvation energy might explain why PROPKA3 energy gives such an interesting result in terms of solubility, although it remains a possibility that PROPKA3 is capturing an insight into solubility which is currently not understood. Pursuing the idea that PROPKA3 energy reflects geometrical changes, on average, as protein length increases, then why is this related to solubility? One possible contributing factor is simply that a longer protein has more centres that can become partially denatured leading to insoluble seeds, but the role of surface geometry suggests an additional possibility.

Incorporated in Fig 7A are the correlations with solubility and length for calculated contact numbers of DEKR groups to AA classes. A scatter plot of solubility and DEKR contacts to aromatic groups within 10 Å (Fig 7D) has a similar form to that of PROPKA3 energy with solubility (Fig 7C). Thus, a further factor to consider is whether partially unfolding regions near the surface of larger proteins carrying a larger complement of hydrophobic groups, due to a less protruding surface, could contribute to the formation of protein-protein interactions. This idea is consistent with the use of chaperone-mediated folding for proteins with more complex topologies in *E. coli* [43] and local partial unfolding in multi-domain proteins observed by hydrogen-deuterium exchange mass spectrometry [44]. With the known role of cation-pi interactions at functional protein interfaces [45], it is reasonable to suggest that they may also play a role in formation of insoluble aggregates.

## 4. Conclusion

Two of the features correlating most highly with solubility in the protein-sol model are [K – R] (positive correlation), and protein length (negative correlation). It was hypothesised that R may be relatively depleted in higher solubility proteins due to a greater ability to form protein-protein interactions, particularly where partial protein unfolding gives greater flexibility for Arg and increased exposure of hydrophobic groups. This model would fit with the increased propensity of Arg to form cation-pi interactions relative to Lys [45]. Considering that the termini of α-helices tend to be relatively exposed to solvent and may be regions of increased propensity for partial unfolding, [K – R] composition was examined throughout helices in protomer AlphaFold models for *E. coli* proteins. Lys composition does indeed increase relative to Arg for the most soluble proteins. This observation adds to the known N-cap and C-cap AA preferences [24], but with a specific connection to protein solubility.

Analysis of [K – R] correlation with solubility for AA classes, helical and non-helical, higher and lower pLDDT, shows that a correlation persists to varying degrees throughout AA classes, and that there are also systematic differences in [K – R] averages between the AA classes, for example with most Lys relative to Arg in lower pLDDT regions. Increased Lys relative to Arg at helical termini, on average, in the most soluble *E. coli* proteins reflects the overall correlation of [K – R] with solubility, but the extent to which Arg could seed insolubility in other protein regions, including those less structurally ordered, could be further investigated experimentally.

An overall increase in overall K,R,D,E composition at helical termini is unsurprising due to the more solvent exposed nature of the termini and is enhanced in the most soluble proteins. Whilst an increase in charge is likely to give rise to increased ionisable group contributions to stability at helical termini, ΔpKa predictions suggest that the interaction per group is mostly not enhanced at helical termini. Overall distributions of ΔpKa predictions for two methods (PROPKA3 and pkcalc) are largely similar, but with a notable difference being a higher number of ionisable groups with mildly destabilising ΔpKas in PROPKA3 relative to pkcalc. This is likely the reason that predicted ionisable group contributions to protein stabilisation have a higher proportion of destabilised proteins for PROPKA3 than pkcalc. If it is accepted that most proteins are net stabilised by ionisable group interactions at pH 7 [42], then this aspect of the PROPKA3 results may arise for ionisable groups with few stabilising charge-charge interactions that are moderately over-balanced by too great a desolvation contribution. This effect is small, in previous comparison of PROPKA3 and pkcalc the PROPKA3 ΔpKas performed better in assessing known pH-dependence in proteins [39].

The mildly destabilising peak of PROPKA3 ΔpKas does, however, lead to an intriguing observation, that PROPKA3 pH 7 energy (per AA) correlates with protein length, and inversely correlates with solubility. Perhaps the PROPKA3 calculation reflects some energetic property that is key for protein solubility, but it is not clear what that might be. A preferred explanation lies with the measures of SASA and surface protrusion that also correlate with protein length. It is proposed that the variation of these geometrical measures with protein length leads to overestimates of the PROPKA3 desolvation term. In terms of solubility, the surface proximal regions in larger proteins with overall less surface protrusion, have larger complements of hydrophobic (aromatic in particular) groups that could be take part in protein-protein interactions, upon partial unfolding.

Studying two of the features that drive the protein-sol model for protein solubility, for [K – R] the role of Arg depletion in more soluble proteins is reinforced, whilst it is suggested that length correlation with solubility could be partly attributable to partial unfolding of regions that include aromatic groups. This work brings together those two features in the context of a potential role for cation-pi interactions in the formation insoluble aggregates.

## Acknowledgements

The authors thank Shalaw Sallah and Robin Curtis for valuable discussions, and staff at the University of Manchester Computational Shared Facility for facilitating storage and processing of data.

## Notes

### Competing Interest Statement

The authors have declared no competing interest.

